# Long-distance dispersal, ice sheet dynamics, and mountaintop isolation underlie the genetic structure of glacier ice worms

**DOI:** 10.1101/463521

**Authors:** Scott Hotaling, Daniel H. Shain, Shirley A. Lang, Robin K. Bagley, Lusha M. Tronstad, David W. Weisrock, Joanna L. Kelley

**Author notes:** **Correspondence:** Scott Hotaling, School of Biological Sciences, Washington State University, Pullman, WA, 99164, USA.

## Abstract

Disentangling the contemporary and historical factors underlying the spatial distributions of species is a central goal of biogeography. For species with broad distributions but little capacity to actively disperse, disconnected geographic distributions highlight the potential influence of passive, long-distance dispersal (LDD) on their evolutionary histories. However, dispersal alone cannot completely account for the biogeography of any species, and other factors–e.g., habitat suitability, life history–must also be considered. North American ice worms (*Mesenchytraeus solifugus*) are ice-obligate annelids that inhabit coastal glaciers from Oregon to Alaska. Previous studies identified a complex biogeographic history for ice worms, with evidence for genetic isolation, unexpectedly close relationships among geographically disjunct lineages, and contemporary migration across large (> 1,500 km) areas of unsuitable habitat. In this study, we analyzed genome-scale sequence data for most of the known ice worm range. We found clear support for divergence between populations along the Pacific Coast and the inland flanks of the Coast Mountains (mean F_ST_ = 0.60), likely precipitated by episodic ice sheet expansion and contraction during the Pleistocene. We also found support for LDD of ice worms from Alaska to Vancouver Island, perhaps mediated by migrating birds. Our results highlight the power of genomic data for disentangling complex biogeographic patterns, including the presence of LDD.

## Introduction

For more than a century, long-distance dispersal (LDD) among presumably isolated populations has intrigued biologists [1–4]. Historically considered rare and unpredictable, the idea that LDD can act as a general mechanism influencing the biogeography of presumably dispersal-limited, macroscopic organisms has gained traction in recent years, with examples accumulating for both plants [5] and invertebrates [6–9]. Many animal vectors play an integral role in plant and invertebrate LDD [e.g., 10, 11], however, in most cases, the resulting LDD is limited to less than 10 km. For more extreme LDD events (e.g., greater than 100 km), the most common animal vector is likely migratory birds, as they seasonally move by the millions over broad spatial scales and geographic barriers, visiting similar habitats along the way [12, 13]. Through this mechanism, dispersal units (e.g., whole organisms, eggs, seeds, etc.) may be ingested and dispersed after passing through the digestive tract [9] or by directly adhering to the bird’s exterior [12]. Thus, as long as there is an opportunity for migratory birds and dispersal units to interact, the opportunity also exists for LDD. Physically quantifying LDD is difficult, however, because it requires real-time sampling and searching (internal and external) of migrating birds for hitchhiking dispersers. Moreover, because rare migratory events can affect species distributions [14] and influence genetic differentiation among populations [15], even thorough physical surveys of migratory birds that find no evidence for LDD cannot rule out its presence. Therefore, alternative approaches for detecting and characterizing LDD should be employed. Because population genomic tools are well-suited to detecting gene flow and genetic structure among populations [e.g., 16], these tools are also well-suited to the indirect detection of LDD, even in the absence of field observations.

Many mechanisms influence genetic relationships among taxa and a range of factors should be considered when attempting to reconstruct biogeographic patterns. For instance, pulses and contractions of glaciers and ice sheets have shaped the evolutionary histories of populations and species throughout Earth’s history [17–20]. These ice sheet dynamics typically affect organisms by separating and reconnecting populations as ice cover changes. However, some species are directly tied to ice sheets [e.g., the meltwater stonefly, 17, 21] and are therefore much more susceptible to ice sheet influence on their evolutionary trajectories. Perhaps no species is more directly tied to ice sheets than glacier ice worms, *Mesenchytraeus solifugus* in North America [22] and *Sinenchytraeus glacialis* in Tibet [23]. The geographic range of *M. solifugus* (hereafter “ice worm”) follows a coastal arc from the Chugach Mountains in southeast Alaska to the Cascade Volcanoes of Washington and Oregon [24]. Ice worms cannot tolerate temperatures more than roughly ±7 °C from freezing and require glacier ice for survival and reproduction [25]. With such unique ecology and physiology, and a dispersal-limited life history, the evolutionary history of ice worms since diverging from conspecifics [26, 27] should be relatively simple with gene flow occurring during glacial periods and isolation (paired with genetic drift) driving divergence among mountaintop-isolated populations during interglacial periods. Natural systems, however, are often more complex than expected and indeed, the evolutionary history of ice worms challenges general expectations of gene flow and evolutionary dynamics in ice-dominated, mountain ecosystems.

Previous genetic studies based on one or two genetic markers identified three ice worm lineages: a “northern” clade in southern Alaska, a “central” clade in southeast Alaska and northern British Columbia, and a “southern” clade ranging over much of British Columbia to southern Oregon [24, 25, 27]. Surprisingly, phylogenetic evidence supported the northern and southern lineages as being most closely related to one another despite the central clade separating them geographically. The most curious aspect of ice worm biogeography, however, has been the repeated discovery of closely related ice worms on glaciers several hundred to thousands of kilometers south of their closest genetic relatives [25, 27; P. Wimberger, unpublished data]. These disjunct northern ice worms co-occurred with, but appeared genetically distinct from, their conspecifics (either central or southern clade ice worms) on the same glaciers. Dial *et al.* [25] laid out three possible explanations for this pattern: wind transport, passerine-mediated dispersal, or a more extensive previous range of the northern clade. While wind transport seems unlikely, the potential for passerine-mediated dispersal is reasonable, particularly in light of other examples of bird-mediated LDD [e.g., 7, 28]. The third scenario, a more extensive distribution of the northern clade with holdover lineages inhabiting the same glacier as more recent colonizers could indeed result in more than one distinct lineage on the same glacier. However, this pattern may only apply to mitochondrial DNA (mtDNA) since mtDNA is maternally inherited and does not recombine. For the nuclear genome, unless strong reproductive isolation exists between the holdover lineages and more recent colonizers, genetic differences would be rapidly homogenized by gene flow and recombination. Assuming no selection against migrants, a reasonable expectation given the similarity of habitat across the ice worm range, the frequency of contemporary versus historical mtDNA haplotypes would therefore depend upon time since introduction, scale of migration (i.e., number of introduced haplotypes), and chance.

In this study, we leveraged a modern population genomic toolkit to add new perspective to the age-old challenge of identifying LDD in wild populations. We also provide new insight into how multiple factors can interact to shape the evolutionary history of species. We hypothesized that the biogeographic history of ice worms stemmed from a confluence of factors: extreme LDD, glacier dynamics, and mountaintop isolation. To test this hypothesis, we generated a genome-wide single nucleotide polymorphism (SNP) data set to answer three specific questions: (1) How do the clades previously diagnosed from a small number of markers hold up to genome-wide scrutiny? (2) What, if any, genomic evidence exists for LDD in ice worms? (3) How do the evolutionary relationships among ice worm populations and genetic clusters align with glacial history in the region [e.g., 29]? Beyond a refined view of ice worm evolution, our study confirms that LDD does occur in ice worms, providing an example of LDD in an annelid and a rare population genomic exploration of the process. Moreover, while considerable evidence details the existence of refugia in the Pacific Northwest (PNW) during the Pleistocene [30], few studies have explored how ice sheet dynamics influenced the evolutionary history of species directly tied to them [e.g., 17]. Our results reveal the profound impact that ice sheet formation during the Pleistocene (∼2.5 million – 11,700 years ago) which flowed episodically from the crest of the Coast Mountains [29] may have had on ice worm evolution, possibly precipitating an ongoing speciation event. Broadly, our findings highlight the power for population genomics to capture contemporary evidence of LDD while also providing biogeographic evidence for reconstructing the glacial history of a region.

## Methods

### Sample collection, library preparation, and SNP calling

During the summer of 2009, ice worms were collected from nine glaciers across most of their geographic range (Figs. 1, S1; Table 1). Samples were stored in > 80% EtOH until DNA was extracted from 59 ice worms using a Qiagen DNEasy Blood and Tissue Kit. Double-digest restriction-site associated DNA (ddRAD) sequencing libraries were prepared following Peterson *et al.* [31] with restriction enzymes EcoRI and NlaIII. During library preparation, samples were divided into two groups and each sample was assigned a unique, variable-length barcode [32] which was incorporated during adapter ligation. Size selection for a 350 bp ± 35 bp window was performed with a Pippen Prep (Sage Science), and both sample groups were subsequently amplified using PCR primers containing a group-specific barcode. The 59-sample library was sequenced on one lane of an Illumina HiSeq4000 at the University of Illinois High-Throughput Sequencing and Genotyping Unit with single-end, 100 bp chemistry.

**Table 1.** Sampling information and summary statistics for all ice worm populations included in this study. Abbreviations: *n* = sample size, π = nucleotide diversity, Het = heterozygosity, F_IS_ = inbreeding coefficient, AK = Alaska, BC = British Columbia, VI = Vancouver Island. π, Het, and F_IS_ were calculated for variable sites only. Asterisks indicate populations included in both Dial *et al.* [27] and this study.

| Population | Latitude, longitude | State/Prov. | Elev. (m) | $n$ | $\pi$ | Het | $F_{IS}$ |
| --- | --- | --- | --- | --- | --- | --- | --- |
| Learnard* (LEA) | 60.806, -148.721 | AK | 624 | 3 | 0.114 | 0.100 | 0.026 |
| Davidson* (DAV) | 59.067, -135.551 | AK | 986 | 15 | 0.078 | 0.065 | 0.037 |
| Treaty* (TRE) | 56.586, -130.151 | BC | 1376 | 8 | 0.026 | 0.033 | -0.012 |
| Bear (BEA) | 56.096, -129.681 | BC | 648 | 6 | 0.039 | 0.046 | -0.013 |
| William Brown* (WIB) | 54.611, -129.129 | BC | 1260 | 6 | 0.056 | 0.057 | 0.000 |
| Jacobson* (JAC) | 52.050, -126.072 | BC | 1249 | 5 | 0.108 | 0.106 | 0.005 |
| White Mantle* (WHM) | 50.795, -125.153 | BC | 1764 | 8 | 0.131 | 0.135 | -0.005 |
| Comox* (COM) | 49.545, -125.355 | BC (VI) | 1881 | 4 | 0.091 | 0.072 | 0.034 |
| Mariner (MAR) | 49.460, -125.764 | BC (VI) | 1754 | 4 | 0.098 | 0.084 | 0.024 |

**Figure 1.**
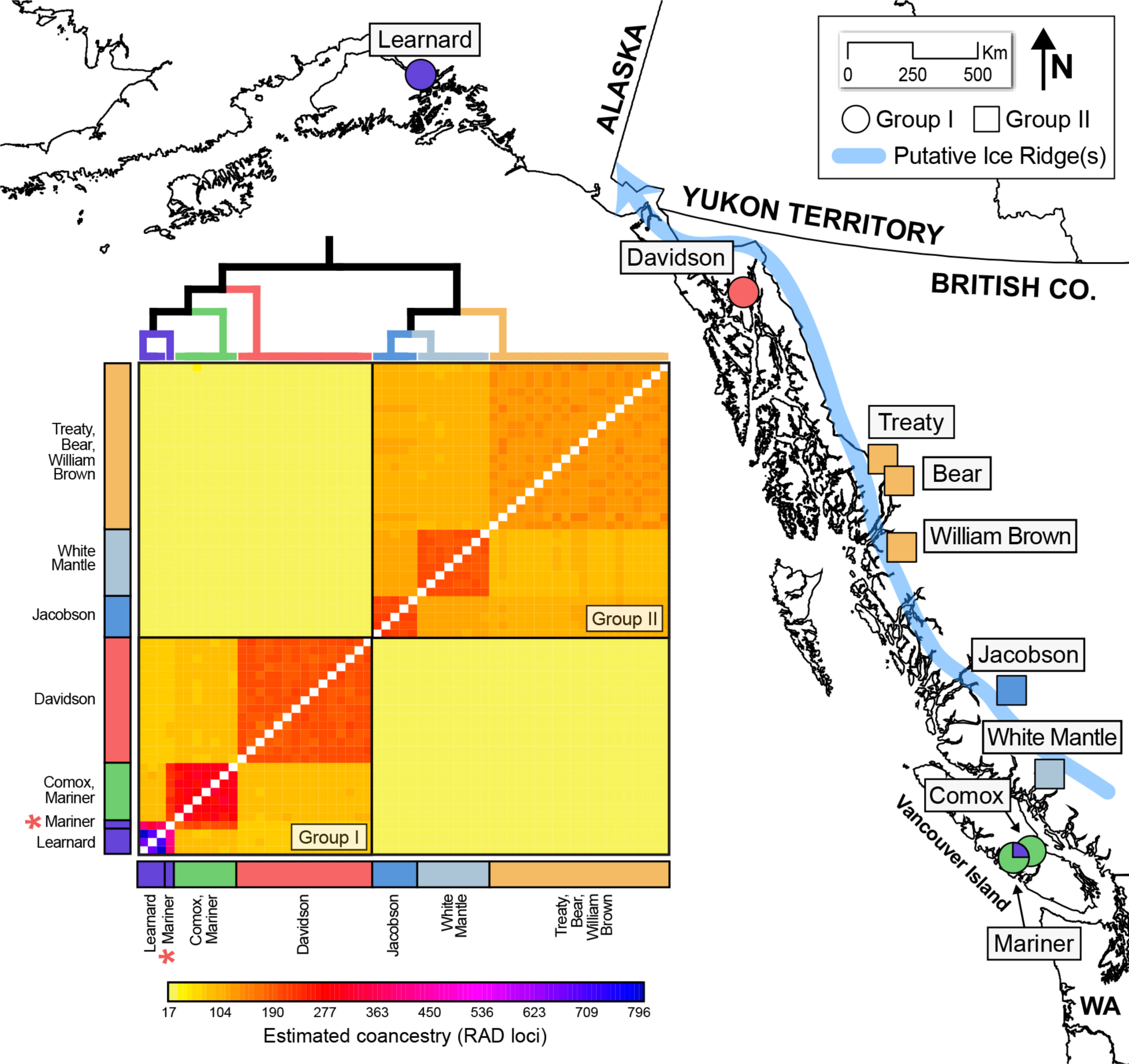
Ice worm populations sampled for this study. Color-coding reflects the results of a fineRADstructure coancestry analysis. Group I (circles) and II (square) populations were generally defined by their presence to the east or west of a key line where ice ridge(s) putatively formed during the Pleistocene > ∼25,000 years ago [29] as well as their distance to the Pacific Ocean. The deep divergence between groups I and II is clearly evident along with more recent differentiation within each group. One individual from the Mariner Glacier (asterisk) was admixed between the Mariner/Comox (Vancouver Island) and Learnard (southern Alaska) clusters, indicating recent LDD.

Raw reads were demultiplexed, quality-filtered, and ddRAD loci were assembled *de novo* using the *process_radtags* and *denovo_map* functions of the Stacks v1.46 pipeline [33]. We allowed a maximum distance between stacks of 2 and a minimum read depth of 10. Next, we applied a stringent filtering scheme to identify high-confidence SNPs that were shared among many individuals. We only included SNPs if they were present in ≥ 5 populations, genotyped in ≥ 50% of individuals per population, and were in Hardy-Weinberg equilibrium with a minor allele frequency of ≥ 0.025 overall. We further restricted analyses to one random SNP per locus for all analyses except fineRADstructure (see below). All post-Stacks filtering steps were performed in PLINK v1.07 [34] and the commands used in this study are provided on GitHub (https://github.com/scotthotaling/ice_worm_ddRAD).

### Population genetic and phylogenetic analyses

For each population, we calculated nucleotide diversity (π), heterozygosity (Het), and the inbreeding coefficient (F_IS_). We also calculated a pair-wise AMOVA F_ST_ for all population combinations [35] with the Stacks *populations* module. To test for a signature of isolation-by-distance [IBD; 36], we estimated Euclidean distances among sites with Google Earth and tested the correlation between geographic distance and F_ST_ with four Mantel tests performed in GenoDive v2.0b27 [37]. The first Mantel test included all nine populations and the second excluded both Vancouver Island populations (Mariner and Comox) to assess whether unique histories for those populations were significantly altering results. The third and fourth Mantel tests focused on signatures of IBD within groups “I” and “II” (see Results). We also measured the Euclidean distance to the Pacific Ocean for each population in Google Earth Pro as an additional spatial comparison of groups I and II. We determined if mean distances to the Pacific Ocean differed between groups with a one-way ANOVA.

Population structure was inferred in two ways: a maximum likelihood-based method using ADMIXTURE 1.3.0 [38] and a discriminant analysis of principal components (DAPC) with the R package *adegenet* [39]. ADMIXTURE analyses were performed with default settings, a range of clusters (*K*) from 1-12, and 25 replicates per *K* with the current time as the random seed. The cross-validation (CV) error for each *K* was plotted to identify the best-fit *K* (minimized CV across the mean of all replicates for each *K*). After identifying the best-fit *K*, we considered the replicate that minimized CV across all 25 replicates for all *K*’s to be the best-fit solution overall. However, because all runs did not converge on the same result, we also inspected best-fit solutions for other replicates of *K* = 7 (the best-fit *K* overall) to clarify the distribution of best-fit solutions. For DAPC, we first used the *find.clusters* function to identify the optimal *K* [i.e., the *K* with the lowest Bayesian Information Criterion (BIC)]. Next, to avoid over-fitting of the model, we retained the appropriate number of principal components (PCs) according to the α-score [PCs retained = 6, Fig. S2]. We performed a final DAPC analysis using the best-fit *K* and optimal number of PCs identified in the previous two steps.

We extended our population structure analyses to infer both shared ancestry and phylogenetic relationships in two ways: a nearest neighbor haplotype approach to infer coancestry with fineRADstructure [40] and phylogenetic relationships inferred from singular value decomposition estimates for quartets of tips using SVDQuartets [41] as implemented in PAUP* v4.0a159 [42]. For fineRADstructure, we used 100,000 burn-in iterations followed by 100,000 iterations sampled every 1,000 steps for the Markov chain Monte Carlo clustering algorithm. Next, we used 10,000 iterations of the tree-building algorithm to assess genetic relationships among clusters. Since fineRADstructure is a haplotype-based approach, analyses were performed using all variable sites for a given ddRAD locus (i.e., a haplotype) rather than randomly selected single SNPs per locus. For SVDQuartets, we performed exhaustive sampling of all possible quartets (every combination of four tips) and branch support was estimated with 100 nonparametric bootstrap replicates.

### Demographic modeling

To test hypotheses of demographic history and estimate divergence times for the two major groups identified in our population genetic and phylogenetic analyses (I and II, see Results), we performed demographic modeling. We used fastsimcoal2 v2.603 [43] which leverages a coalescent-based model to estimate demography from the site frequency spectrum (SFS). We designed and tested the fit of four two-lineage models (Fig. 2A): no gene flow (M1), unidirectional gene flow from group I into II (M2), unidirectional gene flow from group II into group I (M3), and bidirectional gene flow (M4). All four models also included parameters for ancestral (*N*_ANC_) and current (*N*_I_ and *N*_II_) effective population sizes as well as divergence time (T_DIV_). We maximized the number of shared SNPs between our focal groups by selecting the four individuals with the least missing data from the same population in each group (group I = Davidson; group II = Treaty). We also only retained loci with no missing data, which yielded 2,714 SNPs across the eight individuals. For each model, we performed 50 replicate runs, each with 100,000 simulations and 100 cycles of a conditional maximization algorithm. We specified the nuclear mutation rate at 3.5E-9 per site per generation, based on the estimate for *Drosophila melanogaster* [44]. To identify the best-fit model, we calculated an Akaike Information Criterion (AIC) score for each model and selected the best model as the one with the lowest AIC relative to the other models. We generated 95% confidence intervals (CIs) by simulating 50 SFS replicates from the best-fit run of the best-fit model. Next, we performed the same 50 replicate analyses described above for each of the 50 newly simulated SFSs.

**Figure 2.**
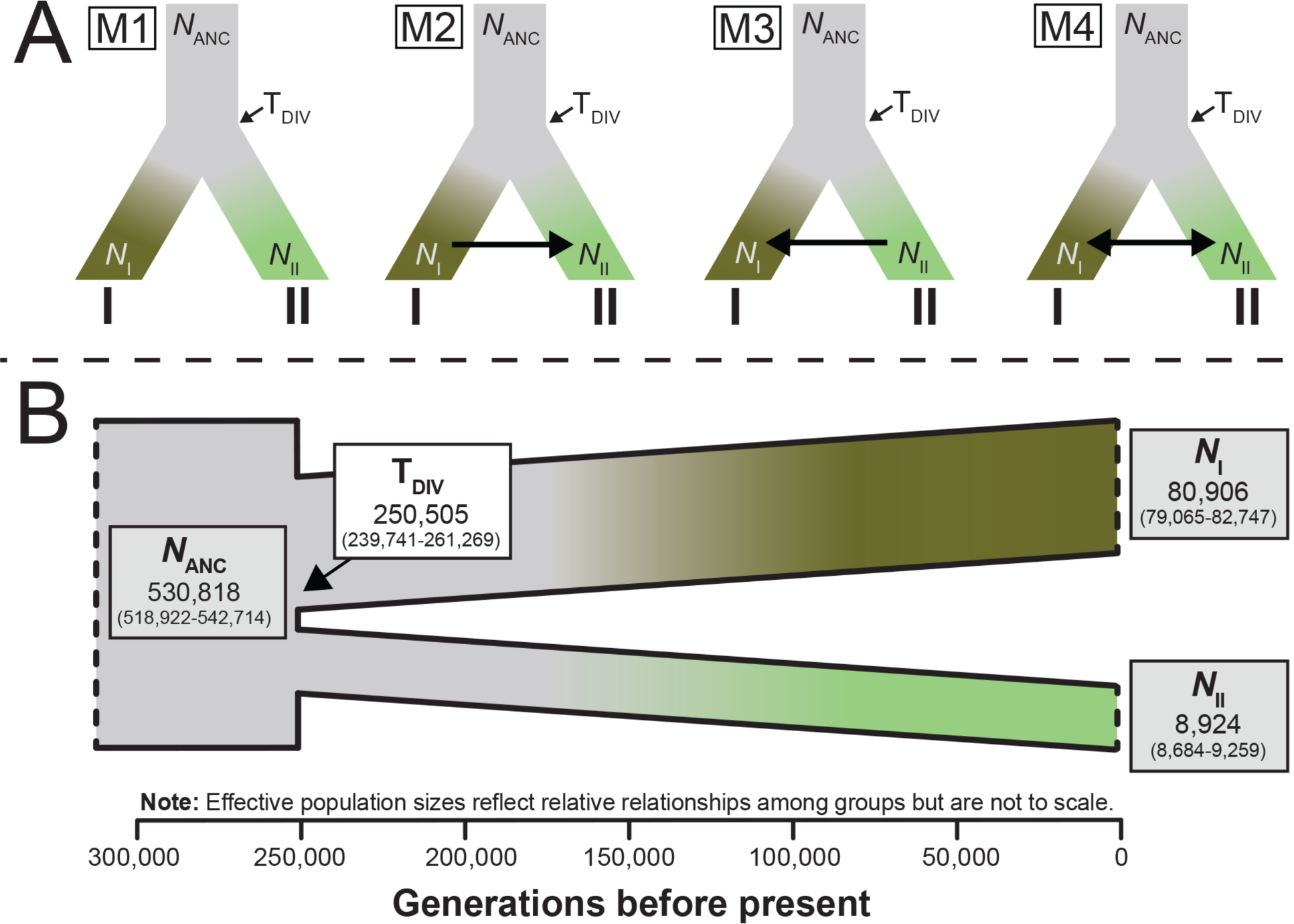
(A) Demographic models tested in this study which included parameters for divergence time (T_DIV_), effective population sizes [ancestral (*N*_ANC_) and for each group (*N*_I_, *N*_II_)], and migration (arrows) when applicable. (B) The parameterized best-fit demographic model (M1). Numbers in parentheses represent 95% confidence interval around estimates. The white box corresponds to the divergence time (in generations) between groups I and II. Gray boxes correspond to population size parameters.

For divergence times, our final results are in generations and we refrain from converting this estimate to years before present because the generation time for ice worms is not known. Estimates of generation time for the family Enchytraeidae also vary widely, ranging from 21 days at 18°C [45] to a full year at 10°C [46]. Moreover, no estimates of generation time at low temperatures (e.g., ∼0°C) are available for the group. Complete details of our demographic analyses are provided in the Supplementary Materials.

## Results

### Population genetic and phylogenetic analyses

We generated 343,875,880 reads with an average of 5,828,404 sequences per individual (min. = 446,872 and max. = 40,982,490). Our total RAD data set included 360,534 unique loci. After filtering, we retained 6,019 loci and 10,392 SNPs (mean = 1.73 SNPs per locus). This final data set had genotype calls for ∼65% of all SNPs. Nucleotide diversity (π) was highest in the White Mantle and Learnard populations (0.131 and 0.114, respectively) and lowest in the Treaty, Bear, and William Brown populations (0.026-0.056; Table 1). Heterozygosity followed the same pattern as π (Table 1). The inbreeding coefficient (F_IS_) was highest in the Davidson and Comox populations (0.037 and 0.034, respectively) and lowest in Bear (−0.013) and Treaty (−0.012; Table 1). Mean differentiation (F_ST_) for all pair-wise comparisons was 0.439. The Learnard population from southern Alaska was, on average, the most differentiated from all others (mean F_ST_ = 0.504) and White Mantle the least differentiated (mean F_ST_ = 0.383; Table 2). We detected no association between genetic and geographic distances in either of the study area-wide Mantel tests (Mantel’s *r*, all populations = −0.04, *P* = 0.42; Mantel’s *r*, no Vancouver Island populations = 0.15, *P* = 0.36). There was, however, a signature of IBD within group II (Mantel’s *r*, group I = 0.839, *P* = 0.025) but not within group I (Mantel’s *r*, group II = 0.839, *P* = 0.082).

**Table 2.** Above the diagonal: Pair-wise AMOVA F_ST_ values for all populations included in this study. Mean F_ST_ (bottom row) refers to the average pair-wise differentiation for columnar populations versus all others. Below the diagonal (in gray): mean pair-wise shared loci for the fineRADstructure coancestry analysis (see Fig. 1).

|  | LEA | DAV | TRE | BEA | WIB | JAC | WHM | COM | MAR |
| --- | --- | --- | --- | --- | --- | --- | --- | --- | --- |
| LEA | -- | 0.333 | 0.634 | 0.626 | 0.608 | 0.586 | 0.557 | 0.347 | 0.344 |
| DAV | 74.9 | -- | 0.567 | 0.557 | 0.541 | 0.527 | 0.515 | 0.290 | 0.295 |
| TRE | 19.8 | 20.0 | -- | 0.153 | 0.207 | 0.275 | 0.250 | 0.656 | 0.652 |
| BEA | 19.0 | 19.9 | 127.2 | -- | 0.165 | 0.250 | 0.222 | 0.650 | 0.645 |
| WIB | 20.1 | 20.5 | 119.8 | 123.0 | -- | 0.216 | 0.201 | 0.630 | 0.628 |
| JAC | 21.0 | 20.4 | 99.7 | 99.5 | 104.7 | -- | 0.172 | 0.604 | 0.600 |
| WHM | 21.6 | 19.1 | 90.7 | 90.9 | 93.0 | 107.8 | -- | 0.576 | 0.572 |
| COM | 84.8 | 86.1 | 21.8 | 21.3 | 23.0 | 22.9 | 22.1 | -- | 0.160 |
| MAR | 164.5 | 82.9 | 21.2 | 21.3 | 22.3 | 21.8 | 21.4 | 262.3 | -- |
| Mean $F_{ST}$ | 0.504 | 0.453 | 0.424 | 0.408 | 0.399 | 0.404 | 0.383 | 0.489 | 0.487 |

Our DAPC analyses supported *K* = 6 as the optimal number of genetic clusters (Fig. 3A,C). ADMIXTURE results, however, supported *K* = 7 as the best-fit (Figs. 3B,C) and the SVDQuartets phylogeny largely mirrored both lines of population structure evidence (Fig. 3D). All analyses supported multiple independent genetic clusters of ice worms. Our DAPC and ADMIXTURE results differed in two ways: (1) the best-fit DAPC result grouped the Treaty, Bear, and William Brown populations into one cluster whereas the best-fit ADMIXTURE result split William Brown into its own cluster. This difference accounted for the *K* = 6 versus *K* = 7 discrepancy between the approaches. (2) While both analyses identified a single individual (MS5) from the Mariner population with genetic assignment to the Learnard cluster, DAPC indicated full assignment of MS5 to the Learnerd cluster whereas Admixture equally assigned it to both the Learnerd and Comox+Mariner clusters (Figs. 1, 3). Finally, because our SNP filtering focused on overarching patterns in the data set, and likely overlooked some degree of population-specific detail, our results are likely conservative estimates of genetic structure in the group.

**Figure 3.**
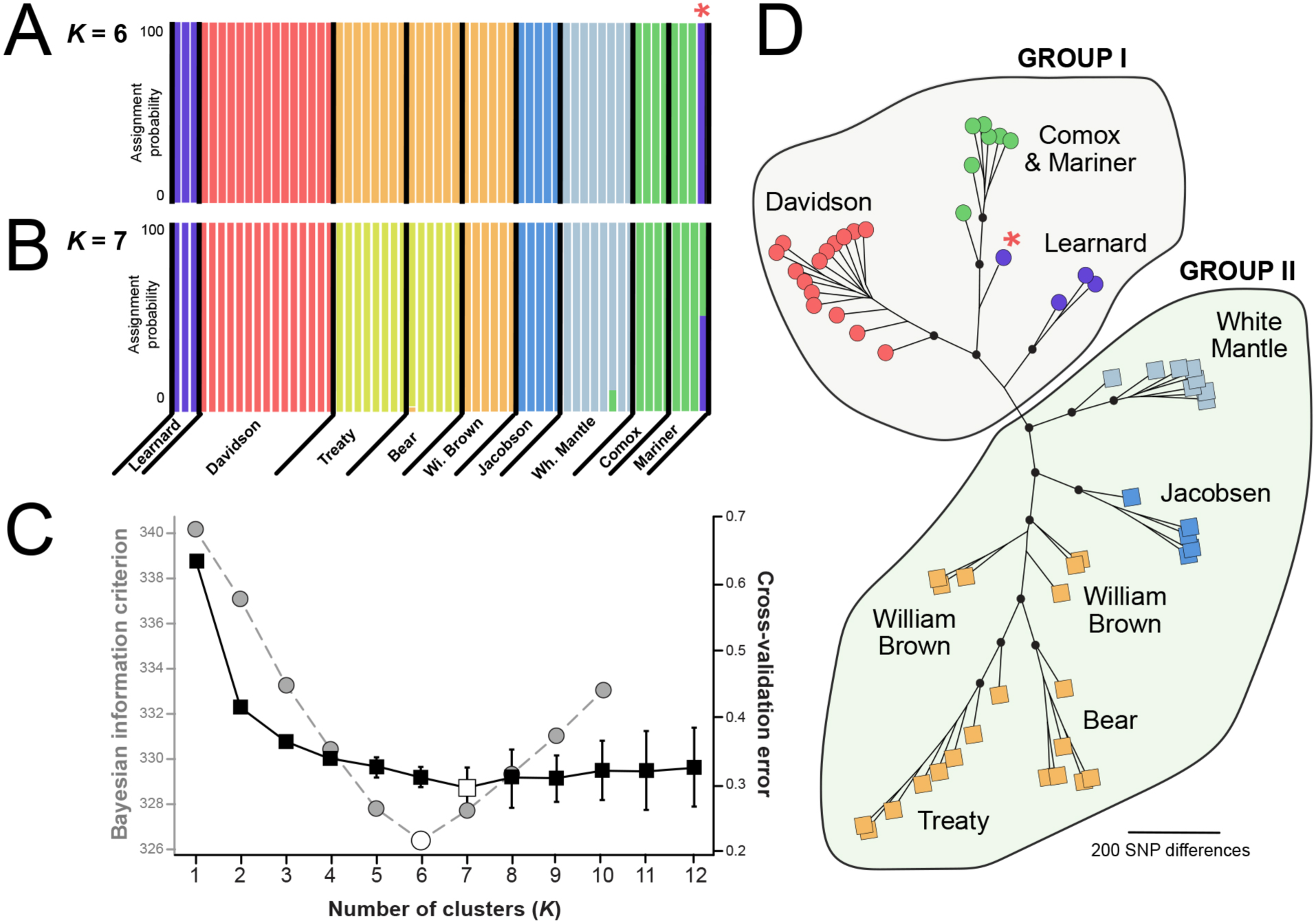
Population genetic structure of ice worms based upon (A) a discriminant analysis of principal components (DAPC) for *K* = 6 and (B) ADMIXTURE results for *K* = 7. (C) Comparisons of support for different values of *K* for DAPC (Bayesian information criterion, BIC; gray dashed line, left y-axis) and Admixture (cross-validation error, CV; dark line, right y-axis). The best-fit *K* (white square, ADMIXTURE; white circle, DAPC) corresponds to the *K* at which CV (ADMIXTURE) or BIC (DAPC) was minimized, respectively. For ADMIXTURE, vertical black bars represent the standard deviation of CV values for each *K* across 25 replicates. (D) An unrooted phylogeny of ice worms generated with SVDQuartets. Tip colorations reflect assignments in (A). Dark circles indicate nodes with greater than 95% bootstrap support. Specimens in group I (circles) and II (squares) are denoted with different symbols. Asterisks (A & D) highlight a single specimen, MS5, which showed evidence of shared ancestry across geographically disjunct populations, indicating LDD.

Our fineRADstructure results largely mirrored those from DAPC, identifying the same six genetic clusters. As in the Admixture results, MS5 exhibited evidence for shared ancestry between the Learnard and Comox/Mariner genetic clusters (Fig. 1). On average, MS5 exhibited a ∼70/30 split of shared ancestry between Learnard and Comox/Mariner ice worms (Fig. 1). Notably, MS5 also exhibited the highest heterozygosity of any individual in the study (and this result did not stem from outsized coverage, Fig. S3). Our fineRADstructure results also highlighted a primary divergence between two groups of ice worm populations (groups I and II; Fig. 1). This split was corroborated by both SVDQuartets (Fig. 3) and F_ST_ comparisons. Mean pairwise F_ST_ among populations within groups I and II were 0.21 and 0.30, respectively. Between groups, however, mean pairwise F_ST_ was 0.60. Mean distance to the Pacific Ocean also differed by 100.5 km (group I = 73.3 km, group II = 173.8 km; ANOVA, *P* < 0.001).

### Demographic modeling

Our tests of demographic models for groups I and II identified a history of divergence without gene flow (model M1) as the best fit to our data (Fig. 2B; Table S1). All other models were rejected with ΔAIC ≥ 9.1 (Table S1). The second-best model (M2) included unidirectional gene flow from group I into group II (Fig. 2A), while models M3 and M4 included gene flow from group II into group I, resulting in ΔAIC scores ≥ 118.4. Groups I and II diverged approximately ∼250,000 generations ago (Fig. 2B).

## Discussion

Historical and contemporary factors, both biotic and abiotic, interact to shape the present-day genetic structure of populations and species. Disentangling their varied contributions can be difficult, however, particularly when evolutionary histories are muddled by unexpected events (e.g., LDD of an organism with limited potential for active dispersal). The modern population genomic toolkit provides historically unprecedented power to resolve biogeographical complexity by allowing more quantitative perspectives of relatedness and greatly improved resolution of genetic independence or similarity [47]. In this study, we used a population genomic data set to refine understanding of the evolutionary history of the extremophile, glacier-obligate ice worm, *M. solifugus*. Our results provide a clear genomic perspective of LDD, showing unequivocally that migration has occurred between southern Alaska and the glaciers of Vancouver Island ∼1,900 km to the south and across the Pacific Ocean. We also provide an independent line of biological evidence in support of the geological hypothesis that ice ridges formed along the crest of the Coast Mountains during the Pleistocene [29].

### Ice worm biogeography and LDD

The recent biogeographic history of ice worms appears to have been shaped by three main factors: i) ice sheet dynamics, ii) mountaintop isolation from conspecifics following the retreat of Pleistocene ice into higher elevations, and iii) LDD.

i. Evidence for the first, overarching factor that has defined the recent evolution of ice worms lies in our overwhelming support for the deepest divergence between two groups (I and II) which fall largely on either side of the Coast Mountains in western North America. During the Pleistocene (∼2.5 million–11,700 years ago), western Canada was repeatedly covered by a continental ice sheet [29]. Ice was generated in the high peaks of the Coast Mountains and subsequently flowed west to the Pacific Ocean and east to the interior of British Columbia from the crest of the range [29, 48, 49]. This potential western-eastern divergence in ice worms aligns with this divide, suggesting that each group diverged in allopatry from their conspecifics. The timeline of this divergence, however, is unclear. Our results suggest ∼250,000 generations have passed since the initial divergence but with no knowledge of generation times for ice worms, nor related species at very low temperatures, we cannot provide a reliable estimate of years before present. If ice worms develop rapidly, with multiple generations per year (e.g., ≥ 3), the divergence may have occurred less than 100,000 years ago. However, if ice worms develop slowly (e.g., one generation for every five years)—the split may have occurred over a million years ago. Either way, the split appears more recent than estimates from mitochondrial DNA of ∼1.7 million years ago [25]. At its maximum, the Pleistocene ice sheet in the region was a ∼2,000-3,000 m high convex dish with gentle interior slopes that steepened at its periphery [29, 50]. Given the sensitivity of ice worms to extreme cold [25], populations likely only persisted on the ice sheet margins, as supported by their present-day occurrences on the lower flanks of higher elevation, low-latitude glaciers. It is possible that, as suggested previously [25, 27], the Boundary Ranges (the most northern subrange of the Coast Mountains), are actually the biogeographic barrier that drove the deep divergence in ice worms described in this study. However, without more fine-scale population genomic sampling on both sides of the Coast Mountains (including the Boundary Ranges), this nuance of ice worm biogeography will remain unclear (see Fig. S4 and additional discussion in Supplementary Materials). In the same vein, the strong genetic similarity of White Mantle to populations east of the proposed ice ridge (Fig. 1), despite falling on its western side, indicates that either the ice ridge that precipitated divergence among the two groups actually formed more to the west than previously thought [29], the White Mantle population has migrated west since divergence from conspecifics, the ice ridge itself was not a barrier driving differentiation (as discussed above), or some combination of all three.
ii. In western North American, the formation of the Cordilleran ice sheet was seeded by alpine glaciers [29]. While the specific dynamics of deglaciation on valley and drainage scales are unknown, a safe assumption is that glaciers retreated from valleys into higher elevations, likely with ice worm populations in tow. Increasing mountaintop isolation and subsequent genetic drift likely precipitated the more recent differentiation within groups I and II since their initial split. Evidence for IBD within group II supports this hypothesis. The lack of support for IBD in group I, however, may reflect reality, the reduced power of a smaller sample size, or LDD maintaining connections over larger spatial scales than a purely IBD model would predict.
iii. Despite limited sampling, we were able to identify one instance of recent LDD among geographically disparate ice worm populations. Indeed, with divided ancestry between the Mariner/Comox and Learnard clusters, one specimen (MS5), is likely the progeny of recent migration between the two. This indicates that LDD is both ongoing in ice worms and perhaps not particularly rare. The most plausible mechanism for ice worm LDD is passive dispersal of mucous-coated ice worm cocoons sticking to the feet, beaks, or feathers of southward migrating birds [12]. Several passerines (e.g., gray-crowned rosy finches, *Leucosticte tephrocotis*) have been observed feeding on ice worms [51, 52; S.H., personal observation] and the presence of an ice worm clitellum [53] indicates that ice worms, like other *Mesenchytraeus* species, reproduce by egg-laden cocoons [25]. The seemingly exclusive northwest-to-southeast pattern of ice worm LDD also has temporal support from bird migratory behavior. Late autumn ice worm reproduction [at the end of the productive season on mountain glaciers, 54] likely occurs in concert with southward-migrating birds stopping to feed on glaciers free of seasonal snow. In contrast, returning spring migrants pass over the same glaciers when seasonal snowfall still covers overwintering ice worms [25], limiting the potential for LDD in the reverse direction.

One question remains, however, if birds are precipitating LDD in ice worms, why has it only been observed for populations west of the Coast Mountains? This curiosity ties in to an important question in North American biogeography: to what extent have ice sheets driven present-day patterns of speciation and genetic differentiation among fauna of the northwest? For ice worms, we hypothesize that populations comprising groups I and II have accumulated some degree of reproductive isolation. This inference is supported by our demographic modeling which rejected models involving gene flow from group II into group I. While this may be at least partly linked to directionality in migration (e.g., bird movements), the strength in which these models were rejected suggests that a zygotic barrier may be limiting inter-group migrants from leaving a genomic signature of gene flow. It is also possible, and perhaps likely, that patterns of LDD in ice worms is driven by vector migration patterns. For instance, *L. tephrocotis*, like other songbirds [55], may preferentially follow coastlines during migration. However, until more is known about the specific interactions of ice worms with various bird species, and by proxy, their potential to act as LDD vectors, relating bird migrations to ice worm distributions and demography will remain difficult. Beyond ice worms, ice sheets have been implicated as a key driver of speciation in boreal birds [56] and phylogeographic structure of many taxa, from nematodes to gray wolves [17, 30, 57–60], and our results clearly support these broader implications for biodiversity accumulation and maintenance in North America.

## Conclusions

In this study, we leveraged population genomic data to unravel the complex evolutionary history of the North American ice worm, *M. solifugus.* Our results add new clarity to previous perspectives on ice worm biogeography while also lending genomic support to the existence of contemporary, likely passerine-mediated LDD in the group. We described two genetic grouping (I and II) which are described with respect to the crest of the Coast Mountains, where ice ridges formed during the Pleistocene [29]. While the phylogenetic data used in this study (i.e., the lack of an outgroup) preclude us from diagnosing groups I and II as monophyletic, given the results of previous studies [24, 25, 27], we predict that future efforts will diagnose them as such, perhaps even representing two nascent species. Finally, our genomic data lend support to the glaciological record in the region, adding a biological line of evidence to a postulated key north-south dividing line along the crest of the Coast Mountains where ice ridges likely formed during the Pleistocene and repeatedly propagated ice flow to the east and west [29]. This potential for genomics to inform the geological record is intriguing and ice worms, as a rare glacier-obligate macroinvertebrate, may be an ideal taxon for similar studies in the future.

## Acknowledgements

We thank Joe Giersch for map-making assistance and the University of Kentucky Center for Computational Sciences as well as the Washington State University Center for Institutional Research Computing for high-performance computational resources. This research was partially funded by NSF award #IOS-082050 to D.H.S.

## Author contributions

S.H., D.H.S., L.M.T, and D.W.W. conceived of and funded the study. S.H., D.H.S., S.A.L, R.K.B., and D.W.W. collected the data. S.H. and J.L.K. analyzed the data and wrote the manuscript with input from D.H.S., R.K.B., and D.W.W. All authors approved the final version.

## Data accessibility

Raw sequence data for this study has been submitted to GenBank under BioProject #PRJNA479335 and code to reproduce the analyses is deposited on GitHub (https://github.com/scotthotaling/ice_worm_ddRAD).

