## Supplementary Materials for "Long-distance dispersal, ice sheet dynamics, and mountaintop isolation underlie the genetic structure of glacier ice worms"

### Methods:

*Demographic modeling*

To explicitly assess the demographic history and timing of divergence for the major groups identified in our population genetic and phylogenetic analyses (I and II; see Results), we performed demographic modeling in fastsimcoal2 v2.603 [1], a coalescent-based program which estimates demography from the site frequency spectrum (SFS). We developed four two-lineage models (Fig. 2A) which included no gene flow (M1), unidirectional gene flow from group I into II (M2), unidirectional gene flow from group II into group I (M3), and bidirectional gene flow (M4). Model definition files (*.est and *.tpl) are provided in Appendix A. To ensure that our models accurately reflected the evolutionary scenario we sought to model, we visualized models in R with the script ParFileInterpreter-v6.3.1.r which is provided with the fastsimcoal2 documentation. Our demographic analyses included an initial set of model selection runs, comparisons of maximum observed and expected likelihoods to select the best-fit model, then parameter estimation for the best-fit model through parametric bootstrapping.

Because demographic inference from the SFS is particularly dependent on the presence of rare alleles and can be biased by missing data [2], we maximized the number of shared SNPs between our focal groups by selecting four individuals from the same population in each group (group I = Davidson; group II = Treaty). We selected Davidson and Treaty because they are the most geographically proximate populations that belong to groups I and II and both populations were robustly sampled for this study. For each population, we selected the four individuals with the least missing data across the same post-filtering data set used in other analyses. Only loci with no missing data were retained for demographic analyses. This yielded 2,714 SNPs across our eight focal individuals. We converted PLINK-formatted allele counts (as output from Stacks) into folded SFS with a modified version of the fs_from_data.py script included in δaδi [2]. After constructing observed SFS files for variable sites, we adjusted the monomorphic counts for all *.obs files to the total number of monomorphic sites for our RAD loci that were 94 bp long (249,064).

For each model, we performed 50 replicate runs with 100,000 simulations and 100 cycles of a conditional maximization algorithm per run. We specified the nuclear mutation rate at 3.5E-9 per site per generation which was previously estimated for *Drosophila melanogaster* [3]. To identify the best-fit model, the maximum expected likelihood (MEL) was compared to the maximum observed likelihood (MOL) for each model replicate. The best-fit run minimized the difference between MEL and MOL for each model's set of fifty replicates. Using these best-fit runs, we calculated an Akaike Information Criterion (AIC) score for each model using the formula: $AIC = [(2k) - (2 \times ln(10) \times MOL)]$, where $k$ is the number of parameters in the model. We identified the best-fit model as the one with the lowest AIC. We calculated the difference between each model AIC and the best-fit model to rank models according to ΔAIC. Use of the AIC allowed for model comparison despite varying numbers of parameters.

To generate 95% confidence intervals (CIs) of parameter estimates for our best-fit model, we used a combination of the best-fit run of the initial model runs and parametric bootstrapping. We first simulated 50 replicates of the SFS from the *.maxL.par file (i.e., the parameter estimates that produced the maximum likelihood) for the best-fit run (minimized difference between MEL and MOL) of the best-fit model (minimized AIC). Next, we performed the same 50 replicate analyses described above for each of the 50 newly simulated SFS files. Finally, we calculated mean parameter estimates and 95% CIs from the 50 best-fit bootstrapping replicates (i.e., the runs with least difference between MEL and MOL for each of the 50 simulated SFSs).

### Discussion:

*Using previous studies to inform the effect of putative ice ridge formation on ice worm evolution*

Our study is the fourth to investigate ice worm biogeography and population genetic structure using molecular data. The first three [4-6] leveraged either 28S or COI data to draw conclusions. In total, 37 ice worm populations (perhaps a few more depending on how disjunct glaciers are grouped) have been sampled for population genetic analyses (Figure S4). Of those, only ~7-10 are within ~100 km of the putative ice ridge and the bulk (26/37 = ~70%) are far to the northwest or southeast of the key area and well within the area of Groups I and II (or previously, the "Northern" and "Southern" clades). Thus, we are unable to further assess our population genomic conclusions with regards to the effects of the putative ice ridge on ice worm structure in light of past genetic data.

**Supplementary Tables:**

**Table S1.** Results of demographic model testing.

| Model | Description | *k* | AIC | ΔAIC | Model choice |
|---|---|---|---|---|---|
| M1 | No gene flow | 4 | 39,278.96 | -- | 1 |
| M2 | Unidirectional gene flow; I → II | 5 | 39,288.06 | 9.10 | 1.E-02 |
| M3 | Unidirectional gene flow; I ← II | 5 | 39,397.36 | 118.40 | 2.E-26 |
| M4 | Bidirectional gene flow; I ↔ II | 6 | 39,397.59 | 118.63 | 2.E-26 |

**Supplementary Figures:**

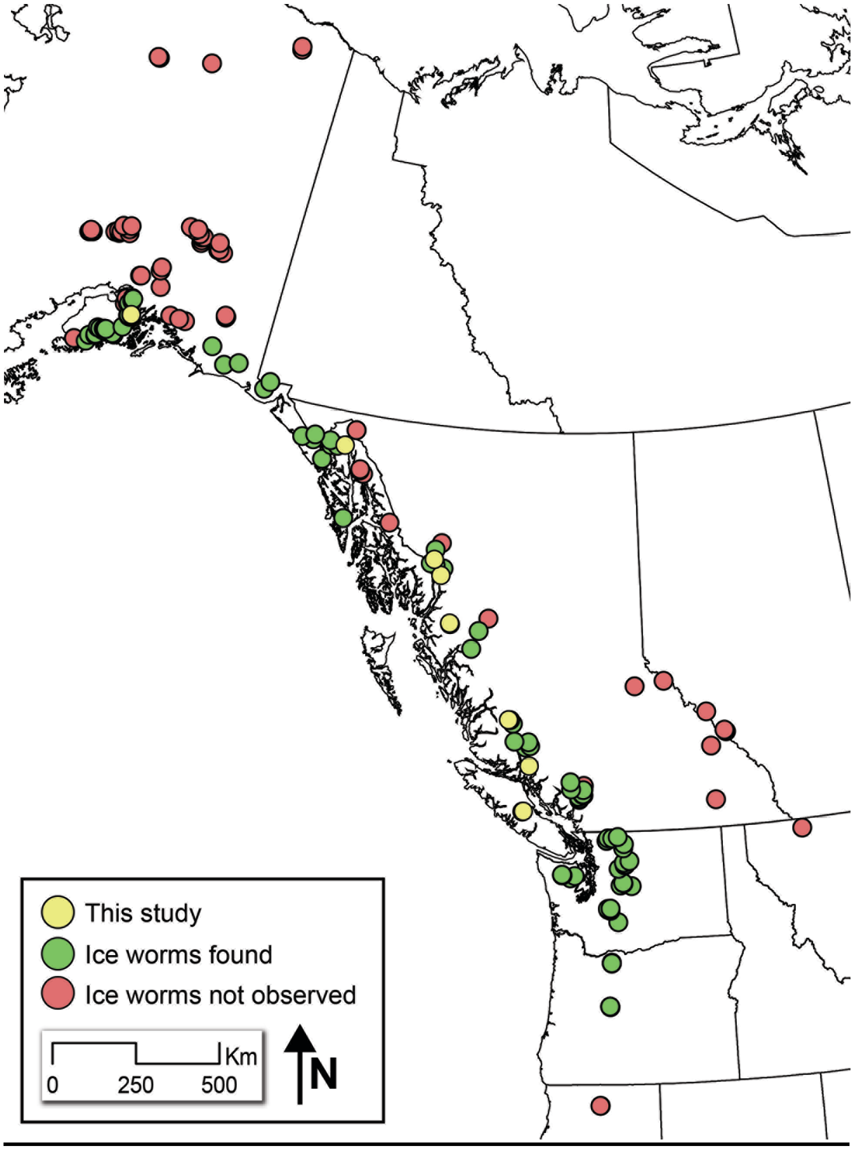


**Figure S1.** The known distribution of ice worms (*Mesenchytraeus solifugus*). Presence and absence information stems from our own surveys, personal communication with Roman Dial, and previous published manuscripts [4-6].

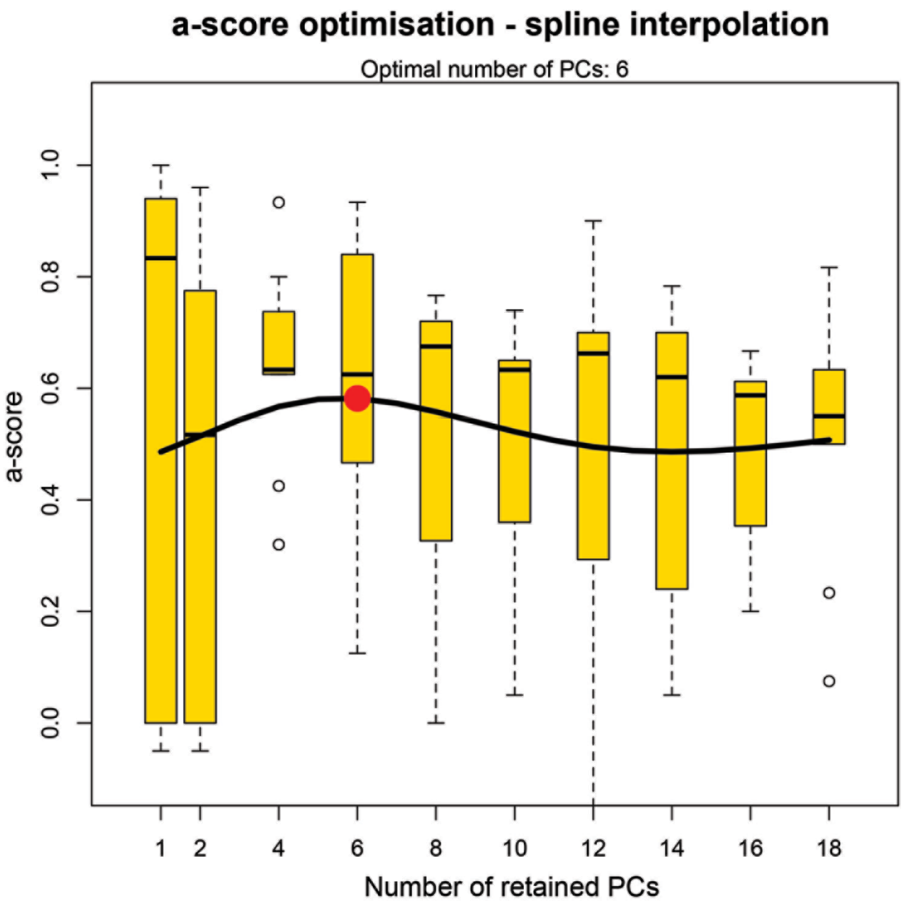


**Figure S2.** An α-score plot for identifying the optimal number of principal components (PCs) to retain in DAPC analyses. The optimal number of PCs to retain (6) is highlighted by a red circle.

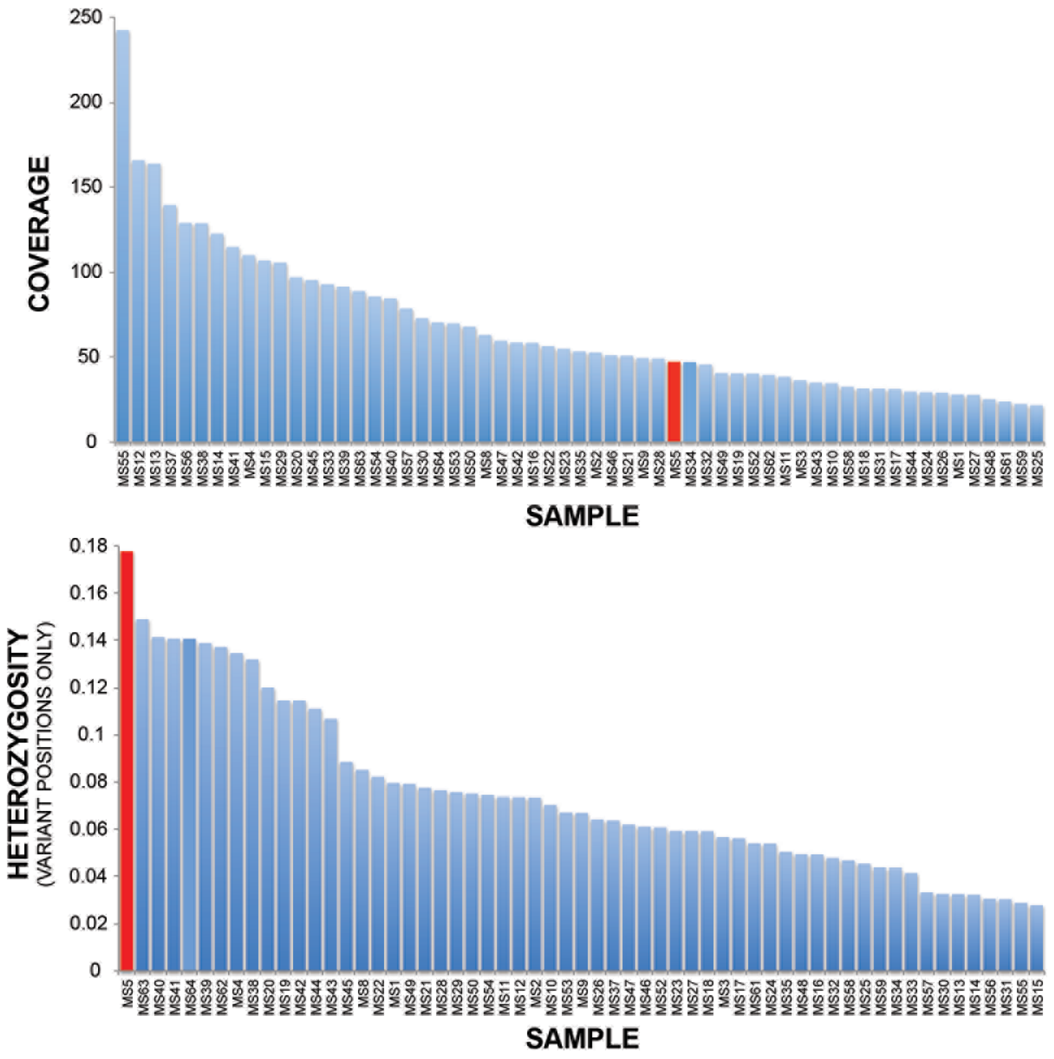


**Figure S3.** Mean coverage and heterozygosity (variant positions only) for all ice worms included in this study. Red bars highlight a single individual (MS5) which bore a signature of substantial admixture between southern Alaska (Learnard Glacier population) and Vancouver Island (Mariner Glacier population).

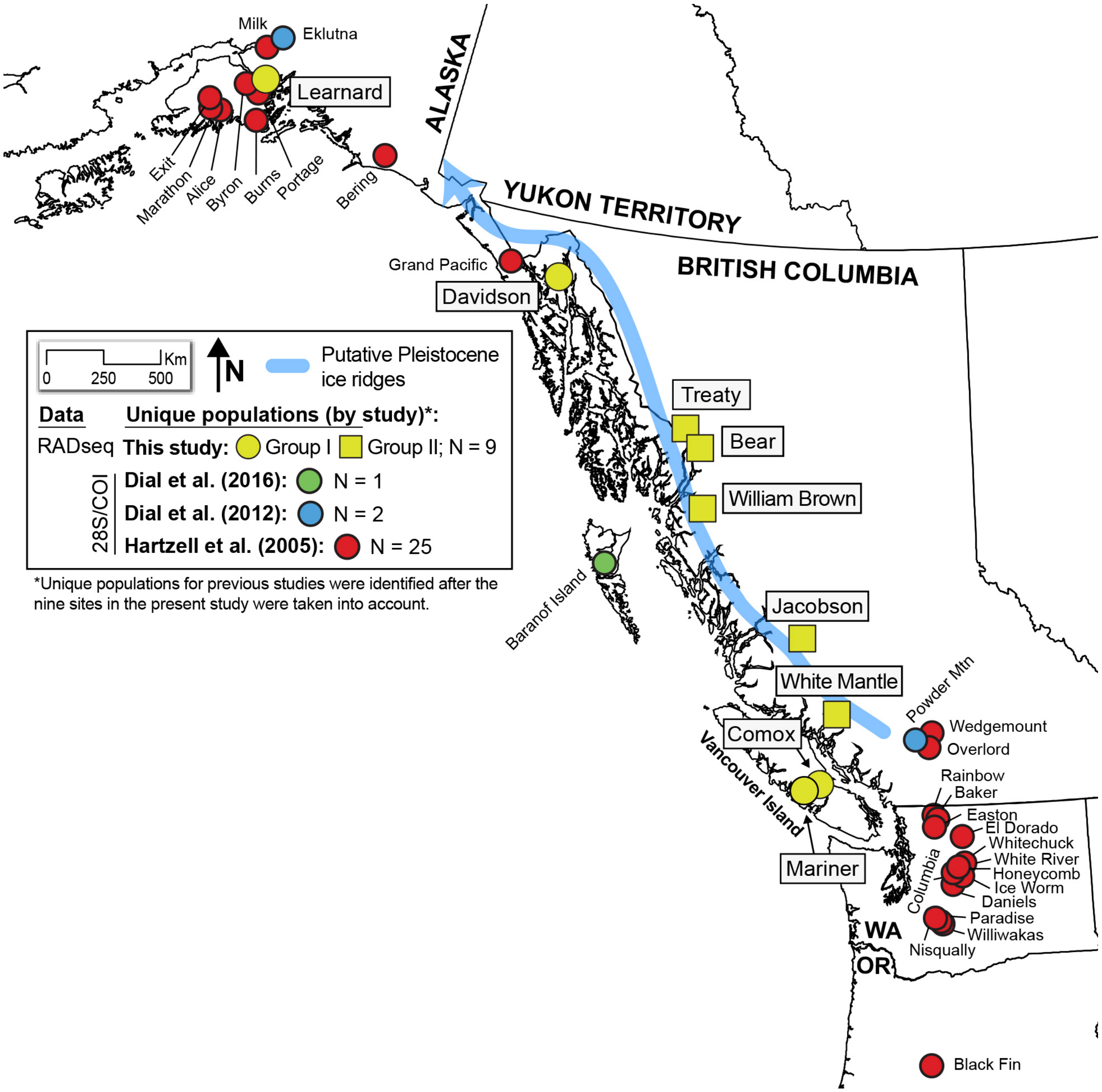


**Figure S4.** All of the sites where population genetic data has been collected for ice worms (*Mesenchytraeus solifugus*). These data include sites from the present study and three previous studies [4-6].

**Appendix A:** fastsimcoal2 model definition files

**M1. est**

```
// Priors and rules file
// ********************

[PARAMETERS]
//#isInt? #name   #dist.#min  #max
//all Ns are in number of haploid individuals
1  ANCSIZE     unif     10    1e6   output
1  NPOP1       unif     10    1e6   output
1  NPOP2       unif     10    1e6   output
1  TDIV        unif     10    1e7   output
[RULES]
[COMPLEX PARAMETERS]
0 RESIZE = ANCSIZE/NPOP1 hide
```

**M1.tpl**

```
//Number of population samples (demes)
2 populations to simulate
//Population effective sizes (number of genes)
NPOP1
NPOP2
//Sample sizes
8
8
//Growth rates
0
0
//Number of migration matrices : 0 implies no migration between demes
2
//Migration matrix 0
0 0
0 0
//Migration matrix 1
0 0
0 0
//historical event: time, source, sink, migrants, new deme size, growth rate, migr mat index
1 historical event
TDIV 1 0 1 RESIZE 0 1
//Number of independent loci [chromosomes]
1 0
//Per chromosome: Number of linkage blocks
1
//per block: Datatype, numm loci, rec rate and mut rate + optional parameters
FREQ 1 0 2.8e-9 OUTEXP
```

**M2. est**

```
// Priors and rules file
// *********************

[PARAMETERS]
//#isInt? #name   #dist.#min  #max
//all Ns are in number of haploid individuals
1  ANCSIZE     unif     10    1e6   output
1  NPOP1       unif     10    1e6   output
1  NPOP2       unif     10    1e6   output
0  N1M21       logunif  1e-2  20    hide
1  TDIV        unif     10    1e7   output
[RULES]
[COMPLEX PARAMETERS]
0 RESIZE = ANCSIZE/NPOP1 hide
0 MIG21 = N1M21/NPOP1 output
```

**M2.tpl**

```
//Number of population samples (demes)
2 populations to simulate
//Population effective sizes (number of genes)
NPOP1
NPOP2
//Sample sizes
8
8
//Growth rates
0
0
//Number of migration matrices : 0 implies no migration between demes
2
//Migration matrix 0
0 MIG21
0 0
//Migration matrix 1
0 0
0 0
//historical event: time, source, sink, migrants, new deme size, growth rate, migr mat index
1 historical event
TDIV 1 0 1 RESIZE 0 1
//Number of independent loci [chromosomes]
1 0
//Per chromosome: Number of linkage blocks
1
//per block: Datatype, numm loci, rec rate and mut rate + optional parameters
FREQ 1 0 2.8e-9 OUTEXP
```

**M3. est**

```
// Priors and rules file
// *********************

[PARAMETERS]
//#isInt? #name   #dist.#min  #max
//all Ns are in number of haploid individuals
1  ANCSIZE     unif     10    1e6   output
1  NPOP1       unif     10    1e6   output
1  NPOP2       unif     10    1e6   output
0  N2M12       logunif  1e-2  20    hide
1  TDIV        unif     10    1e7   output
[RULES]
[COMPLEX PARAMETERS]
0 RESIZE = ANCSIZE/NPOP1 hide
0 MIG12 = N2M12/NPOP2 output
```

**M3.tpl**

```
//Number of population samples (demes)
2 populations to simulate
//Population effective sizes (number of genes)
NPOP1
NPOP2
//Sample sizes
8
8
//Growth rates
0
0
//Number of migration matrices : 0 implies no migration between demes
2
//Migration matrix 0
0 0
MIG12 0
//Migration matrix 1
0 0
0 0
//historical event: time, source, sink, migrants, new deme size, growth rate, migr mat index
1 historical event
TDIV 1 0 1 RESIZE 0 1
//Number of independent loci [chromosomes]
1 0
//Per chromosome: Number of linkage blocks
1
//per block: Datatype, numm loci, rec rate and mut rate + optional parameters
FREQ 1 0 2.8e-9 OUTEXP
```

**M4. est**

```
// Priors and rules file
// *********************

[PARAMETERS]
//#isInt? #name   #dist.#min  #max
//all Ns are in number of haploid individuals
1  ANCSIZE     unif     10    1e6   output
1  NPOP1       unif     10    1e6   output
1  NPOP2       unif     10    1e6   output
0  N1M21       logunif  1e-2  20    hide
0  N2M12       logunif  1e-2  20    hide
1  TDIV        unif     10    1e7   output
[RULES]
[COMPLEX PARAMETERS]
0 RESIZE = ANCSIZE/NPOP1 hide
0 MIG21 = N1M21/NPOP1 output
0 MIG12 = N2M12/NPOP2 output
```

**M4.tpl**

```
//Number of population samples (demes)
2 populations to simulate
//Population effective sizes (number of genes)
NPOP1
NPOP2
//Sample sizes
8
8
//Growth rates
0
0
//Number of migration matrices : 0 implies no migration between demes
2
//Migration matrix 0
0 MIG21
MIG12 0
//Migration matrix 1
0 0
0 0
//historical event: time, source, sink, migrants, new deme size, growth rate, migr mat index
1 historical event
TDIV 1 0 1 RESIZE 0 1
//Number of independent loci [chromosomes]
1 0
//Per chromosome: Number of linkage blocks
1
//per block: Datatype, numm loci, rec rate and mut rate + optional parameters
FREQ 1 0 2.8e-9 OUTEXP
```
